# *Pseudomonas aeruginosa* Stimulates Inflammation and Enhances KSHV-Induced Cell Proliferation and Cellular Transformation through Both LPS and Flagellin

**DOI:** 10.1101/2020.10.07.330803

**Authors:** Ashley Markazi, Paige M. Bracci, Michael McGrath, Shou-Jiang Gao

## Abstract

Inflammation triggered by innate immunity promotes carcinogenesis in cancer. Kaposi’s sarcoma (KS), a hyperproliferative and inflammatory tumor caused by Kaposi’s sarcoma-associated herpesvirus (KSHV) infection, is the most common cancer in AIDS patients. KSHV infection sensitizes cells to pathogen-associated molecular patterns (PAMPs). We examined the role of *Pseudomonas aeruginosa* (PA), an opportunistic bacterium that can affect AIDS patients, in inflammation and cell proliferation of KSHV-transformed cells. PA stimulation increased cell proliferation and efficiency of colony formation in softagar of KSHV-transformed rat primary mesenchymal precursor (KMM) cells but had no significant effect on the untransformed (MM) cells. PA stimulation also increased cell proliferation of KSHV-infected human B-cells, Bjab, but not the uninfected cells. Mechanistically, PA stimulation resulted in increased inflammatory cytokines and activation of p38, ERK1/2, and JNK mitogen-activated protein kinase (MAPK) pathways in KMM cells while having no obvious effect on MM cells. PA induction of inflammation and MAPKs were observed with and without inhibition of Toll-like receptor 4 (TLR4) pathway while a flagellin-deleted mutant of PA required a functional TLR4 pathway to induce inflammation and MAPKs. Furthermore, treatment with both LPS or flagellin alone was sufficient to induce inflammatory cytokines, activate MAPKs, and increase cell proliferation and efficiency of colony formation in softagar of KMM cells. These results demonstrate that both LPS and flagellin are PAMPs that contribute to PA induction of inflammation in KSHV-transformed cells. Because AIDS-KS patients are susceptible to PA infection, our work highlights the preventive and therapeutic potential of targeting PA infection in these patients.

**Importance:** Kaposi’s sarcoma (KS), caused by infection of Kaposi’s sarcoma-associated herpesvirus (KSHV), is one of the most common cancers in AIDS patients. KS is a highly inflammatory tumor but how KSHV infection induces inflammation remains unclear. We have previously shown that KSHV infection upregulates Toll-like receptor 4 (TLR4), sensitizing cells to lipopolysaccharide (LPS) and *Escherichia coli*. In the current study, we examined the role of *Pseudomonas aeruginosa* (PA), an opportunistic bacterium that can affect AIDS patients, in inflammation and cell proliferation of KSHV-transformed cells. PA stimulation increased cell proliferation, inflammatory cytokines, and activation of growth and survival pathways in KSHV-transformed cells through two pathogen-associated molecular patterns LPS and flagellin. Because AIDS-KS patients are susceptible to PA infection, our work highlights the preventive and therapeutic potential of targeting PA infection in these patients.

## Introduction

Inflammation is the immune system’s response to tissue damage or microbial infection. When unregulated, inflammation can cause or exacerbate carcinogenesis (1). For example, the continuous induction of inflammatory cytokines and cytotoxic mediators such as reactive oxygen or nitrogen species can over time cause damage to the cellular genome, resulting in cellular mutations that lead to dysregulated cell proliferation and cancer development (2). Inflammatory cytokines can further enhance cancer progression by activating pro-oncogenic and -survival cellular pathways including NF-κB and STAT3 pathways (3). By further elucidating the mechanisms by which inflammation promotes cancer cell growth, proliferation and survival, we can gain important insights into the mechanism of oncogenesis and identify novel therapeutic targets and biomarkers.

Interestingly, patients with HIV/AIDS are at a much higher risk for developing AIDS-associated cancer (AAC) compared to those with healthy immune systems (4). HIV patients are 500 times more likely to develop Kaposi’s sarcoma (KS), 12 times more likely to be diagnosed with non-Hodgkin’s lymphoma, and, among women, 3 times more likely to be diagnosed with cervical cancer (4). Moreover, HIV patients are at a higher risk for non-AIDS-defining cancer (NADC) as well, including anal cancer, liver cancer, lung cancer, and Hodgkin’s lymphoma (5). HIV enhances cellular transformation through its protein, Tat, which represses tumor suppressor gene p53, promotes cell cycle progression, and induces inflammation (6, 7). For cancers caused by infections of oncogenic viruses, HIV regulates both the replication of these viruses and the progression their associated cancers (8). For examples, HIV-encoded products Tat, Nef, and Vpr regulate KSHV replication and the functions of KSHV genes resulting in enhanced cell migration, invasion, and angiogenesis (9-15). Moreover, long term use of antiretroviral drugs in HIV patients is associated with an elevated risk of several cancers (16).

A number of other mechanisms also contribute to the chronic inflammation and increased cancer risk in HIV-infected patients. One of these factors is immunodeficiency associated with HIV infection such as declined CD4+ T cell count (17). Because CD4+ T cells stimulate B cells and CD8+ T cells, patients with a low CD4+ T cell count are less capable of eliminating infections including viruses, bacteria, and fungi, hence increasing pathogen-associated molecular patterns (PAMPs) (18), which continuously stimulate immune and cancer cells to secrete inflammatory cytokines, resulting in chronic inflammation. Indeed, many reports show that a lower CD4+ cell count is correlated with an increase in inflammatory cytokines (19). Moreover, chronic stress such as metabolic stress of the immune and cancer cells induced by HIV infection, as well as the long-term use of antiretroviral drugs, triggers damage-associated molecular patterns (DAMPs), which further promote chronic inflammation in the same fashion as PAMPs (20).

KS, the most common cancer in HIV-infected patients, is a hyperproliferative and inflammatory cancer caused by infection with Kaposi’s sarcoma-associated herpesvirus (KSHV) (21). KSHV infection provides an excellent model for examining the complex interactions of HIV, a cancer-causing virus (KSHV), innate immunity, inflammation, and cancer. KSHV encodes numerous genes that directly contribute to cellular transformation (21) and KSHV infection alone is sufficient to induce inflammatory cytokines, which can stimulate cell proliferation and survival and regulate KSHV replication (22-28). Furthermore, we have shown that KSHV infection sensitizes the infected cells to PAMPs, leading to the activation of Toll-like receptor 4 (TLR4) and alternative complement pathways, which induce inflammatory cytokines, promote cell survival, proliferation and cellular transformation (22, 29). *Escherichia coli*- and lipopolysaccharide (LPS)-activated TLR4 pathway stimulates cell proliferation, cellular transformation, and tumorigenesis by increasing IL-6 expression to activate the STAT3 pathway (22). Indeed, in a recent study, we have shown the impoverishment of oral microbial diversity and enrichment of specific microbiota in oral KS in HIV-infected patients (30). However, the precise mechanism how specific microbiota promote KS remains to be elucidated.

*Pseudomonas aeruginosa* (PA) is normally considered as a commensal bacterium. However, PA can cause severe infection in individuals with immunosuppression (31). HIV/AIDS patients with CD4+ T cell counts below 200 cell/mm^3^ are at a significantly higher risk for PA infection (32). PA consists of PAMPs, such as LPS and flagellin that activate TLR4 and TLR5, respectively (33). Hence, PA infection might induce inflammatory cytokines of KSHV-infected cells and promote cell proliferation and cellular transformation.

In the current study, we analyzed the effects of PA on cell proliferation and cellular transformation in a KS-like model of KSHV-induced cellular transformation of rat primary embryonic metanephric mesenchymal precursor cells (MM) (34). We observed that PA stimulation increased both cell proliferation and cellular transformation in KSHV-transformed MM cells (KMM) yet had no significant effect on MM cells. Moreover, we observed similar results of increased cell proliferation in a KSHV-infected human B cell line, KSHV-BJAB, compared to the BJAB uninfected control. In KMM cells, PA stimulation resulted in increased expression of inflammatory cytokines and activation of p38, ERK1/2, and JNK pathways. Interestingly, we observed induction of inflammatory cytokines and activation of the p38 and ERK1/2 pathways even after inhibition of TLR4 pathway in KMM cells stimulated by PA, and that this effect disappeared when KMM cells were stimulated with a flagellin-deleted mutant of PA, indicating that PA mediated inflammation and cellular transformation of KSHV-transformed cells through both LPS and flagellin.

## Results

### PA stimulation enhances cell proliferation and cellular transformation of KMM cells but has no significant effect on MM cells

To examine the effect of PA on the proliferation of KSHV-transformed cells, we treated the cells with 1×10^7^ CFU/mL PA (ATCC^®^ 15442™) or 1 μg/mL LPS. PA increased the proliferation of KMM cells but did not have any significant effect on MM cells (Fig. 1A). Similar results were observed with LPS as expected (22). Both PA and LPS also increased the sizes and efficiency of colony formation in softagar of KMM cells (Fig. 1B and C). As previously reported, MM cells did not form any significant colonies (34). These results indicated that similar to LPS, PA stimulated the proliferation and cellular transformation of KMM cells (22). To assess the effects of PA and LPS stimulation on KSHV-infected human B cells, we treated BJAB and KSHV-BJAB cells with 1×10^7^ CFU/mL PA (ATCC^®^ 15442™) or 1 μg/mL LPS. Although less pronounced than in KMM cells, PA stimulation also increased proliferation of KSHV-BJAB cells while having no significant effects in BJAB cells (Fig. 1D). Because KMM cells can form colonies in softagar permitting the evaluation of the transforming potential of the cells, we chose to further examine the effect of PA on KMM cells and the control MM cells in subsequent experiments (34).

**FIG 1.**
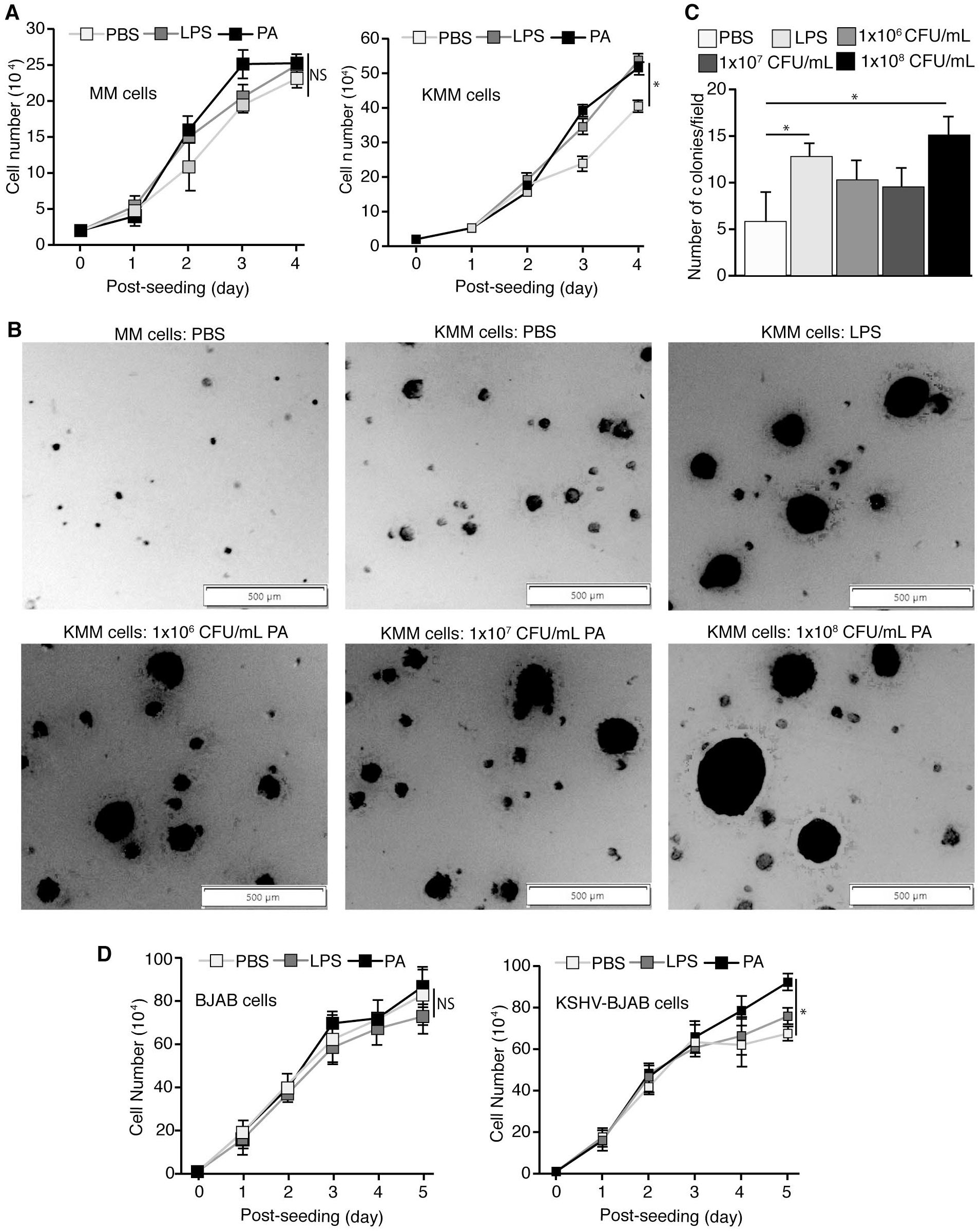
PA stimulation enhances cell proliferation and cellular transformation of KSHV-infected cells but has no significant effect on the uninfected cells. (A) Cell proliferation of MM and KMM cells treated with PBS, 1 μg/mL LPS, or 1×10^7^ CFU/mL PA (ATCC^®^ 15442™) analyzed by cell counting. (B-C) Formation of colonies of KMM cells in softagar treated with PBS, 1 μg/mL LPS, or 1×10^6^-1×10^8^ CFU/mL PA (ATCC^®^ 15442™) shown by representative pictures (B) and results of statistical analysis from 3 wells, each with 5 representative fields (C). (D) Cell proliferation of BJAB and KSHV-BJAB cells treated with PBS, 1 μg/mL LPS, or 1×10^7^ CFU/mL PA (ATCC^®^ 15442™) analyzed by cell counting. Asterisk indicates P ≤ 0.05. NS denotes “not significant”.

### PA stimulation increases the expression levels of inflammatory cytokines in KMM cells while having minimal effect on MM cells

We previously showed that purified *E. coli* LPS induced inflammatory cytokines IL-6, IL-1β, and IL-18 in KMM cells but only had a weak effect on MM cells. PA (ATCC^®^ 15442™) stimulation resulted in higher mRNA levels of IL-6 and IL-1β but had no significant effect on IL-18 in KMM cells (Fig. 2A). Additionally, we analyzed the cytokines TNF-α and CXCL-1 as PA increased levels of these inflammatory cytokines in mice (35, 36). PA stimulation also resulted in higher mRNA levels of TNF-α and CXCL-1 in KMM cells (Fig. 2A). In contrast, cytokines IL-6, IL-1β, IL-18, TNF-α, and CXCL-1 were not significantly upregulated in MM cells by PA (Fig. 2A).

**FIG 2.**
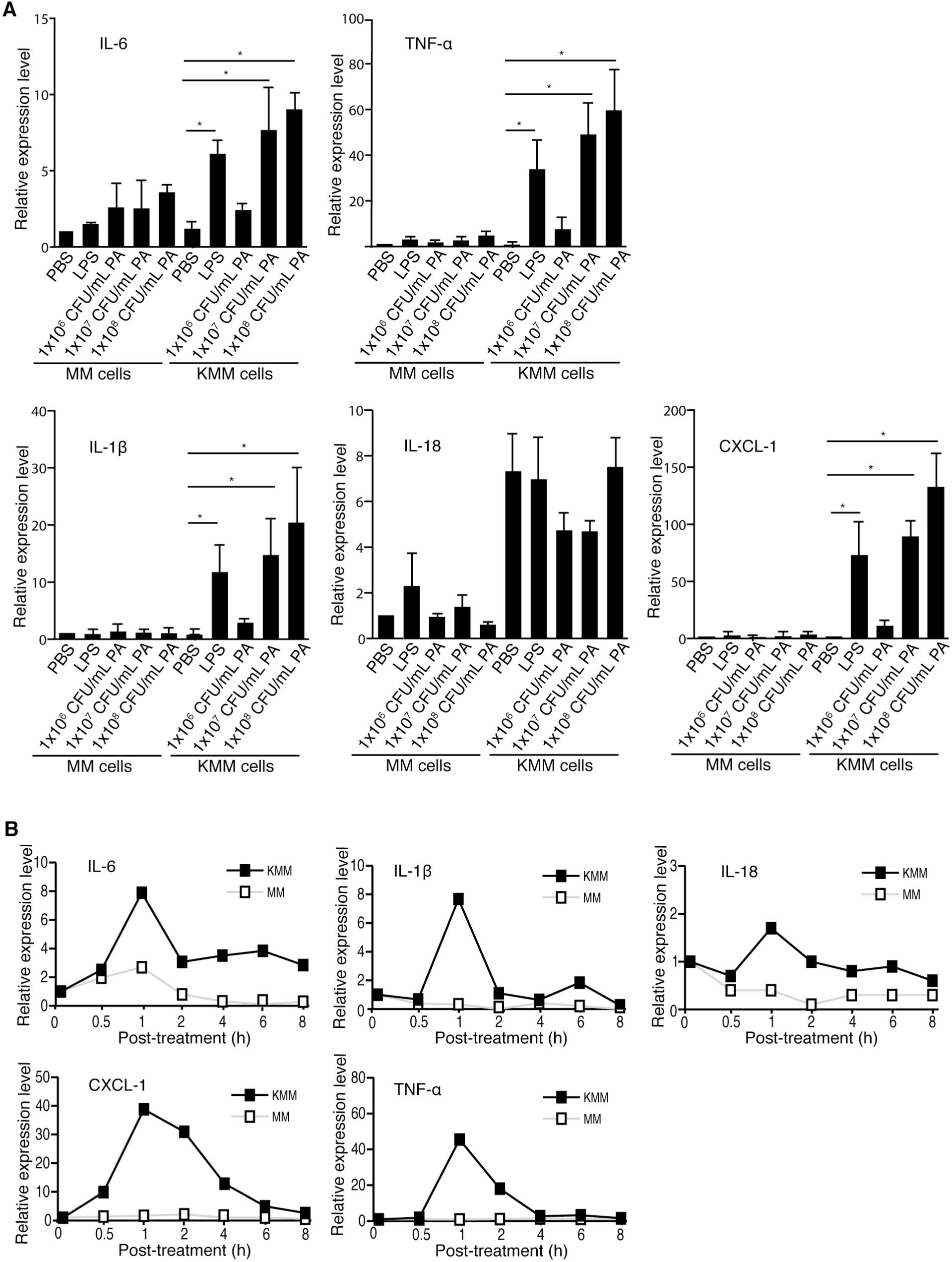
PA stimulation increases the expression of inflammatory cytokines in KMM cells while having minimal effect on MM cells. (A) MM and KMM cells were treated with PBS, 1 μg/mL LPS, or 1×10^6^-1×10^8^ CFU/mL PA (ATCC^®^ 15442™) for 1 h, and analyzed for cytokines IL-6, TNF-α, IL-1β, IL-18, and CXCL-1 by RT-qPCR. (B) MM and KMM cells were treated with PBS or 1×10^7^ CFU/mL PA (ATCC^®^ 15442™) at the specified time points and analyzed for cytokines IL-6, TNF-α, IL-1β, IL-18, and CXCL-1 by RT-qPCR. Asterisk indicates P ≤ 0.05.

We further examined the induction kinetics of inflammatory cytokines in KMM cells by PA by stimulating the cells with 1×10^7^ CFU/mL PA (ATCC^®^ 15442™) and analyzed at 0, 0.5, 1, 2, 4, 6, and 8 h post-stimulation. The IL-6, IL-1β, TNF-α, and CXCL-1 had the highest mRNA levels at 1 h after PA stimulation in KMM cells (Fig. 2B). No significant induction of inflammatory cytokines in different time points in MM cells was observed following PA stimulation (Fig. 2B).

### PA stimulation activates the mitogen-activated protein kinase (MAPK) pathways in KMM cells but has no obvious effect in MM cells

The p38, ERK1/2, and JNK MAPK pathways have been implicated in the induction of inflammatory cytokines and are commonly activated in cancer cells (37). We examined the activation of these pathways following stimulation with 1×10^7^ CFU/mL PA (ATCC^®^ 15442™) in MM and KMM cells. We detected the activation of p38, ERK1/2 and JNK pathways, which peaked at 15 min after PA stimulation in KMM cells (Fig. 3). In contrast, no increased activation of p38, ERK1/2, or JNK was observed in MM cells (Fig. 3).

**FIG 3.**
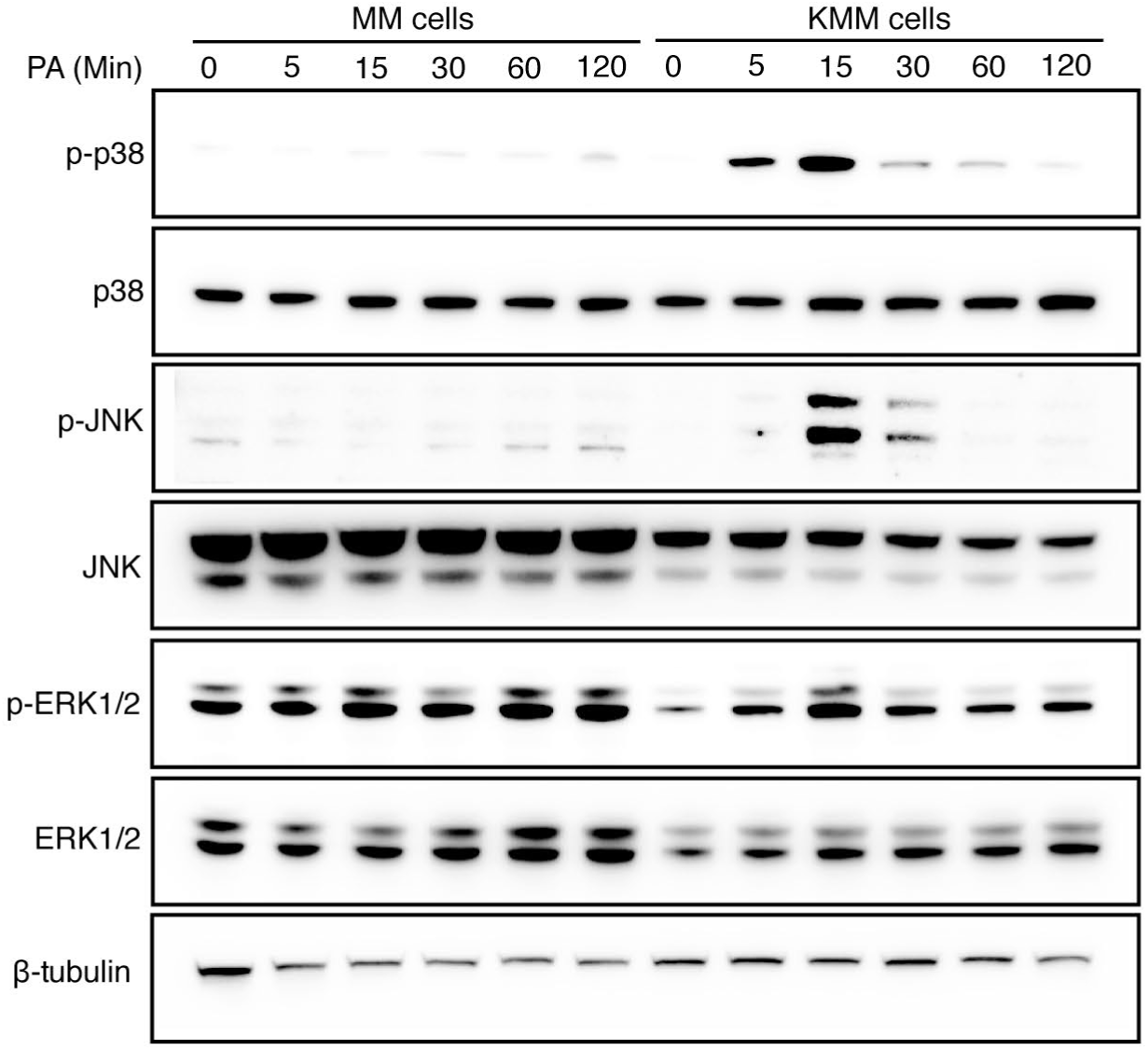
PA stimulation activates the p38, JNK and ERK1/2 MAPK pathways in KMM cells but has no obvious effect on MM cells. MM and KMM cells were stimulated with 1×10^7^ CFU/mL PA (ATCC^®^ 15442™) for the specific time points and analyzed by Western-blotting.

### PA stimulation increases the expression of inflammatory cytokines by both TLR4-dependent and -independent mechanisms in KSHV-transformed cells

We have previously shown that LPS can induce inflammatory cytokines in KMM cells. Since PA contains other PAMPs in addition to LPS, to determine if other PA PAMPs also contributed to the PA-induced inflammation in KMM cells, we stimulated KMM cells with 1×10^7^ CFU/mL PA (ATCC^®^ 15442™) in the presence of 10 μg/mL TLR4 inhibitor CLI095. The levels of induced pro-inflammatory cytokines IL-6, IL-1β, TNF-α, and CXCL-1 by PA were significantly decreased in KMM cells by CLI095; however, they remained significantly higher than the unstimulated KMM cells (Fig. 4). As expected, induction of inflammatory cytokines by LPS was completely blocked by CLI095 in KMM cells with the levels similar to the unstimulated cells or cells treated with CLI095 alone (Fig. 4). No obvious change of inflammatory cytokines in MM cells was observed under these treatments. These results indicated that, besides LPS, other PA PAMP(s) might also contribute to PA-induced inflammation in KMM cells.

**FIG 4.**
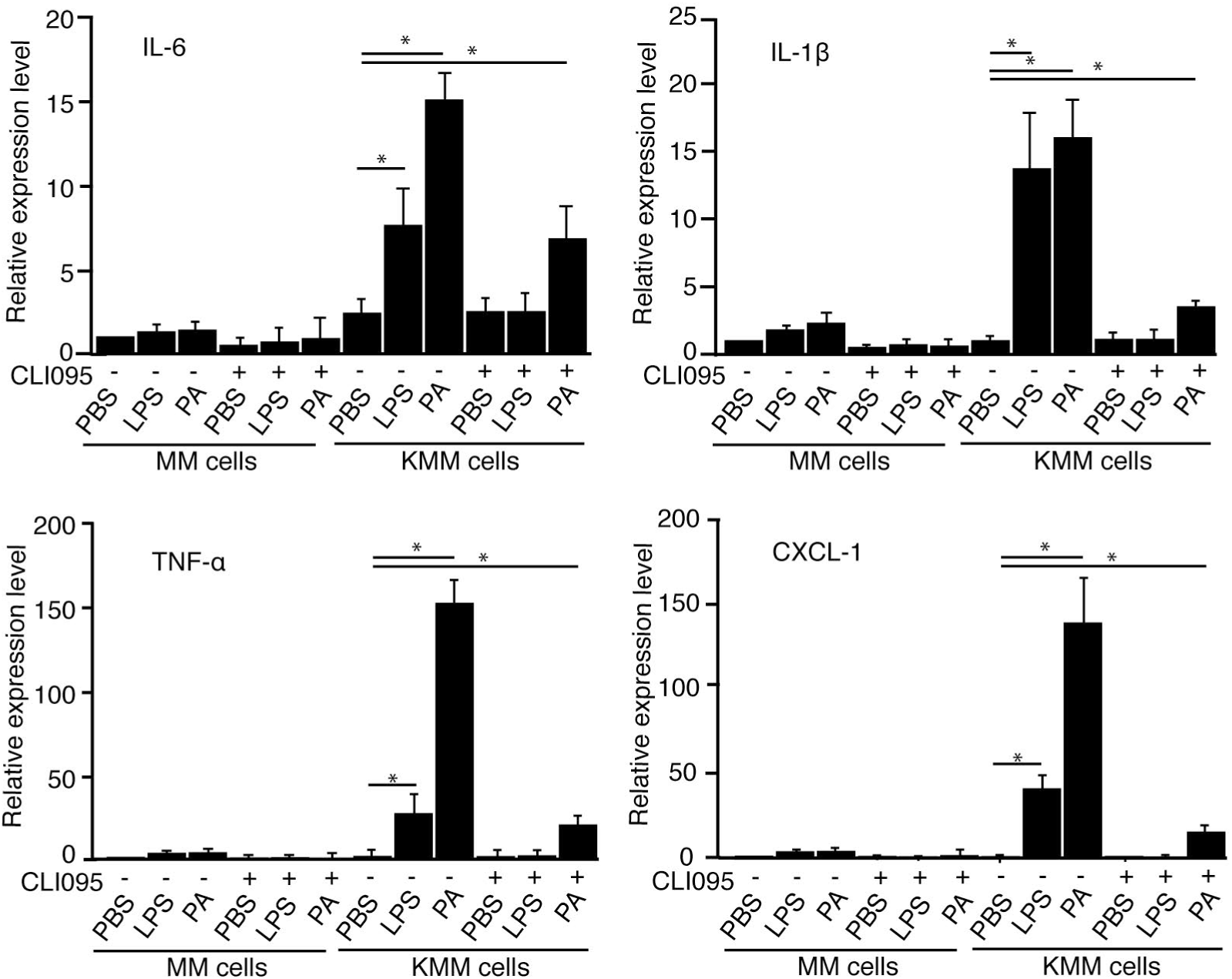
PA stimulation increases the expression of inflammatory cytokines by both TLR4-dependent and -independent mechanisms in KSHV-transformed cells. MM and KMM cells were treated with 10 μg/mL TLR4 inhibitor (CLI095) for 1 h and then treated with PBS, 1 μg/mL LPS, or 1×10^7^ CFU/mL PA (ATCC^®^ 15442™) for 1 h, and analyzed for the expression of cytokines by RT-qPCR. Asterisk indicates P ≤ 0.05.

### PA flagellin contributes to PA induction of inflammatory cytokines in KMM cells but has no significant effect on MM cells

As flagellin is PA’s second most immunogenic PAMP, we stimulated KMM cells with a PA strain with the *fliC* gene deleted from its genome, PAO1Δ*fliC*, which rendered it defective in flagellin expression, and its parallel wild-type PA PAO1 (38). While PAO1 and PAO1Δ*fliC* at 1×10^7^ CFU/mL induced inflammatory cytokines in KMM cells, the levels of induction were significantly lower in KMM cells stimulated with PAO1Δ*fliC* than PAO1 (Fig. 5A). As expected, 1×10^7^ CFU/mL PAO1 induced inflammatory cytokines at levels similar to 1×10^7^ CFU/mL PA (ATCC^®^ 15442™) (Fig. 5A). We then treated MM and KMM cells with CLI095 for 1 h before stimulating with PAO1, PAO1Δ*fliC*, or LPS. Similar to PA (ATCC^®^ 15442™), the PAO1 induction of inflammatory cytokines was reduced in KMM cells by TLR4 inhibitor CLI095 but remained at significant higher levels than the unstimulated cells (Fig. 5B). In contrast, PAO1Δ*fliC* induction of inflammatory cytokines was completely abolished by CLI095 (Fig. 5B). No significant effect was observed on MM cells with either PAO1Δ*fliC* or PAO1 (Fig. 5B).

**FIG 5.**
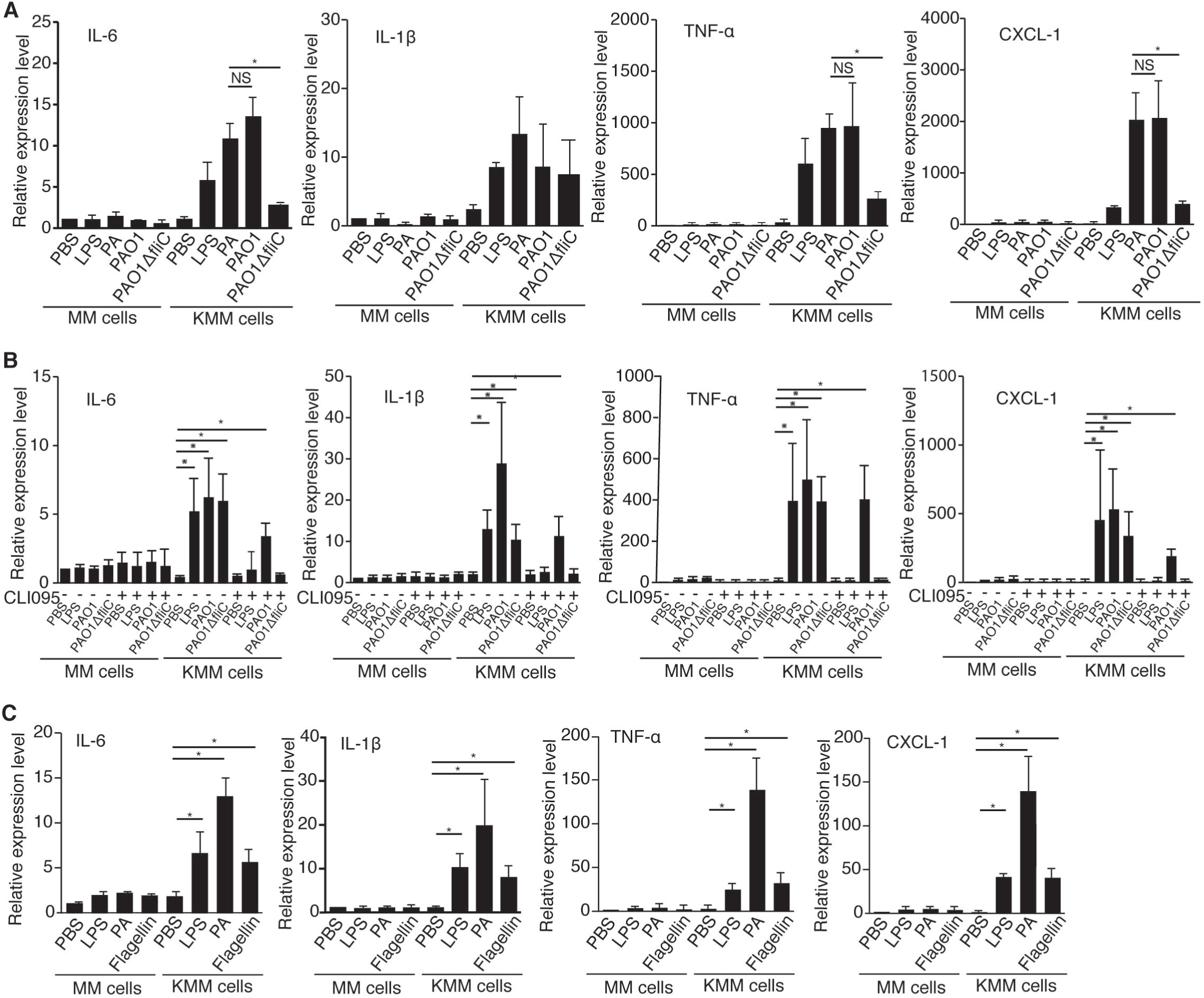
PA flagellin contributes to PA induction of inflammatory cytokines in KMM cells but has no significant effect on MM cells. (A) MM and KMM cells were treated with PBS, 1 μg/mL LPS, 1×10^7^ CFU/mL PA (ATCC^®^ 15442™), 1×10^7^ CFU/mL PAO1, or 1×10^7^ CFU PAO1Δ*fliC* for 1 h, and analyzed for the expression of cytokines by RT-qPCR. (B) MM and KMM cells were treated with 10 μg/mL TLR4 inhibitor (CLI095) for 1 h, and then treated with PBS, 1×10^7^ CFU/mL wild-type PAO1, or 1×10^7^ CFU/mL PAO1Δ*fliC* for 1 h, and analyzed for the expression of cytokines by RT-qPCR. C, MM and KMM cells were treated with PBS, 1 μg/mL LPS, 1×10^7^ CFU/mL PA (ATCC^®^ 15442™), or 0.3 μg/mL PA flagellin for 1 h, and analyzed for the expression of cytokines by RT-qPCR. Asterisk indicates P ≤ 0.05. NS denotes “not significant”.

To further confirm the role of flagellin in PA induction of inflammation in KMM cells, we stimulated the cells with 0.3 μg/mL purified PA flagellin. Treatment with purified PA flagellin alone was sufficient to induce inflammatory cytokines in KMM cells at levels similar to LPS alone (Fig. 5C). In contrast, purified PA flagellin stimulation had no significant effect on MM cells (Fig. 5C).

Taken together, these results indicated that flagellin contributed to PA induction of inflammatory cytokines in KMM cells. Hence, at least LPS and flagellin contributed to PA induction of inflammation in KMM cells.

### PA flagellin activates p38 and ERK1/2 pathways in KMM cells but has no obvious effect on MM cells

The above results indicated that the patterns of inflammatory cytokines induced by LPS and flagellin were not entirely the same, which could be due to distinct activation of the signaling pathways by the two PAMPs. While all three MAPK pathways were activated in KMM cells by PAO1, the activation levels were reduced in cells stimulated with PAO1Δ*fliC* (Fig. 6A). Treatment with TLR4 inhibitor CLI095 reduced PAO1 activation of p38 and ERK1/2 pathways but completely abolished that of JNK pathway, and PAO1Δ*flliC* activation of all three MAPK pathways (Fig. 6A). These results indicated that LPS mediated PA activation of all three pathways while flagellin mediated the activation of p38 and ERK1/2 pathways. We then investigated whether flagellin alone was sufficient to activate the p38 and ERK1/2 pathways in KMM cells, by treating the cells with purified PA flagellin. As expected, flagellin alone indeed activated the p38 and ERK1/2 pathways, and treatment with CLI095 had no effect on flagellin activation of these two pathways albeit the reduction of ERK1/2 activation by CLI095 in PA-stimulated KMM cells was less pronounced in this experiment indicating possible variation of PA LPS effect on ERK1/2 activation (Fig. 6B). Together, these results indicated that flagellin contributed to PA activation of p38 and ERK1/2 pathways, and LPS contributed to PA activation of all three MAPK pathways in KMM cells.

**FIG 6.**
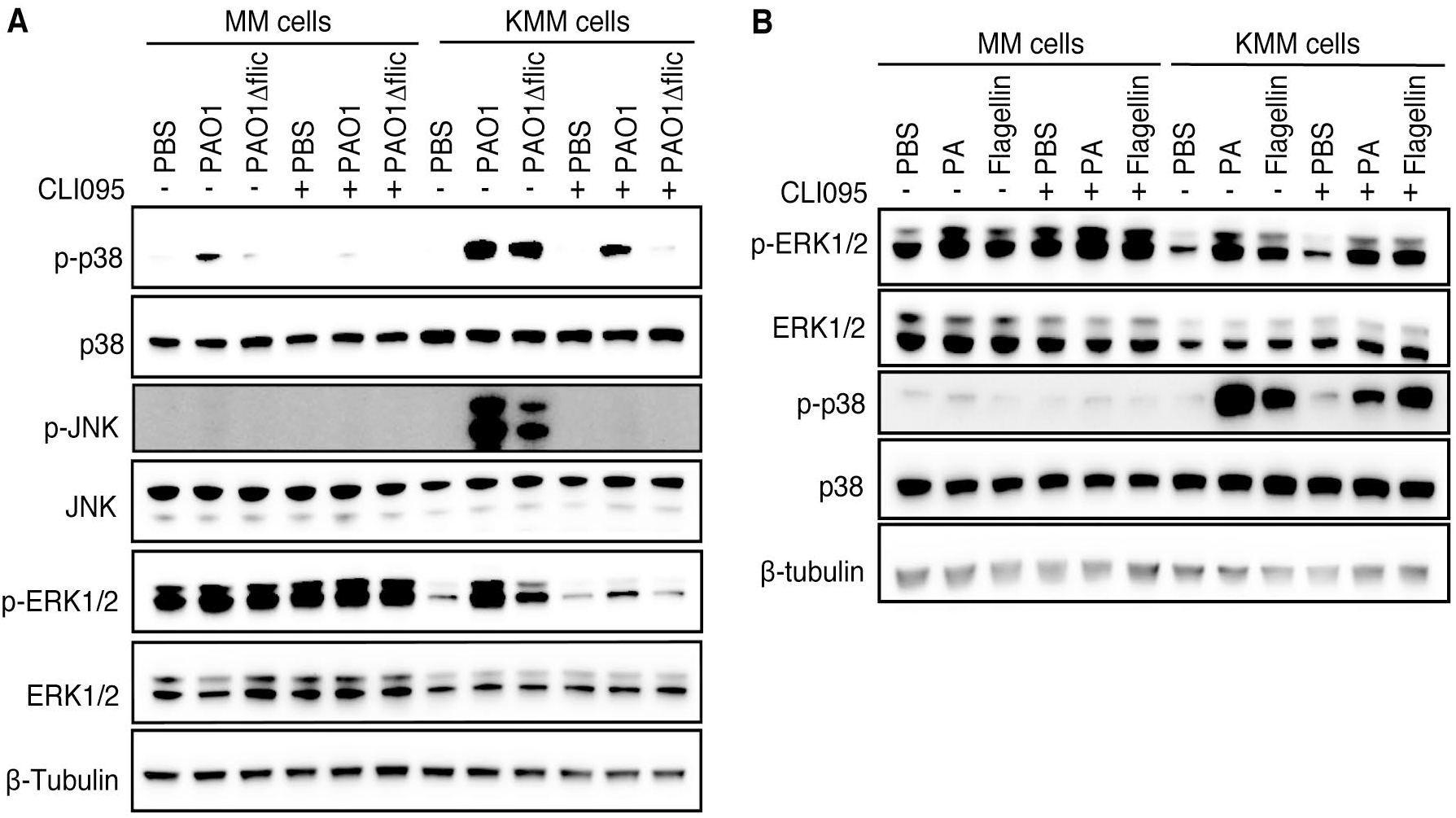
PA-flagellin induces p38 and ERK1/2 activation in KMM cells but has no obvious effect on MM cells. (A) MM and KMM cells were treated for 1 h with 10 μg/mL TLR4 inhibitor CLI095 and then treated with PBS, 1×10^7^ CFU/mL wild-type PAO1, or 1×10^7^ CFU/mL PAO1Δ*fliC* for 15 min, and analyzed for the activation of p38, ERK1/2, and JNK pathways by Western-blotting. (B) MM and KMM cells were treated for 1 h with 10 μg/mL TLR4 inhibitor CLI095 and then treated for 15 min with PBS, 1×10^7^ CFU/mL PA, or 0.3 μg/mL PA-flagellin, and then analyzed for the activation of p38 and ERK1/2 pathways by Western-blotting.

### PA flagellin stimulation increases cell proliferation and cellular transformation of KMM cells but has no significant effect on MM cells

Because flagellin participated PA-induced inflammation and activation of MAPK pathways, we examined whether flagellin could promote KSHV-induced cell proliferation and cellular transformation. KMM cells were treated with purified PA flagellin or PBS as control and analyzed for cell proliferation. Treatment with PA flagellin alone was sufficient to induce faster cell proliferation in KMM cells than the untreated cells (Fig. 7A). In contrast, PA flagellin had no effect on the proliferation of MM cells (Fig. 7A).

**FIG 7.**
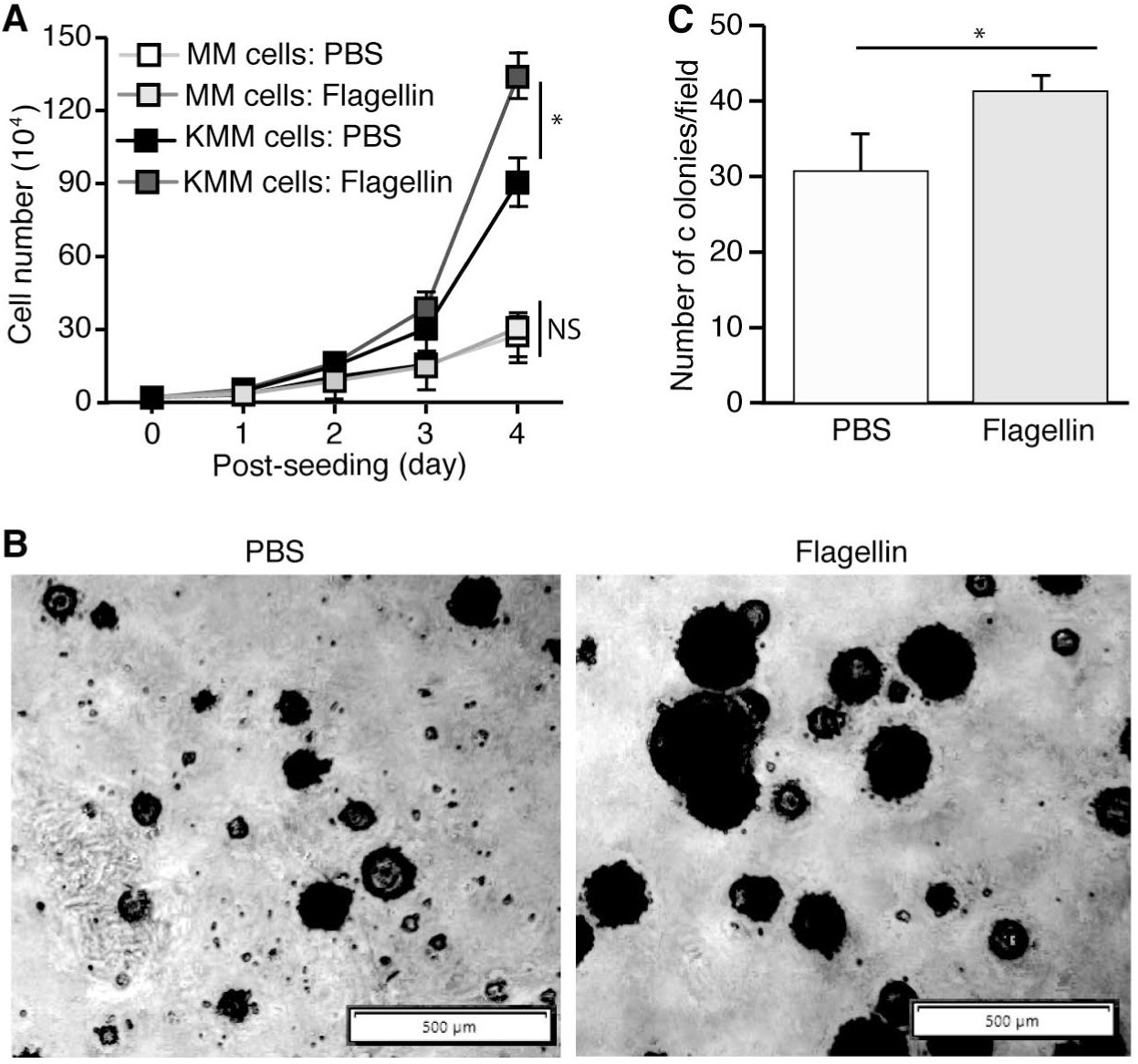
PA flagellin stimulation increases cell proliferation and cellular transformation of KMM cells but has no significant effect on MM cells. (A) Cell proliferation of MM and KMM cells treated with PBS or 1 μg/mL PA-flagellin. (B-C) Formation of colonies of KMM cells in softagar treated with PBS or 1 μg/mL PA-flagellin shown by representative pictures (B) and results of statistical analysis from 3 wells, each with 5 representative fields (C). Asterisk indicates P ≤ 0.05. NS denotes “not significant”.

We further examined the effect of flagellin on cellular transformation of KMM cells. Treatment with purified PA flagellin increased the number and sizes of colonies of KMM cells compared to the untreated cells (Fig. 7B and C). MM cells did not form any observable colonies in softagar in both flagellin treated and untreated cells (results not shown).

### Simultaneous inhibition of p38 and ERK1/2 pathways decreases flagellin-induced inflammation and cell proliferation of KMM cells but has no obvious effect on MM cells

To determine whether p38 and ERK1/2 pathways mediated flagellin-induced inflammation in KMM cells, we stimulate the cells with PA flagellin in the presence of specific inhibitors of these pathways. p38 inhibitor (SB 203580) or ERK1/2 inhibitor (U0126) alone had no effect on flagellin-induction of inflammatory cytokines (Fig. 8A). However, treatment with both inhibitors significantly decreased the flagellin induction of all inflammatory cytokines (Fig. 8A), confirming the important roles of p38 and ERK1/2 pathways in flagellin induction of inflammatory cytokines. Hence, both p38 and ERK1/2 pathways mediated flagellin induction of inflammatory cytokines.

**FIG 8.**
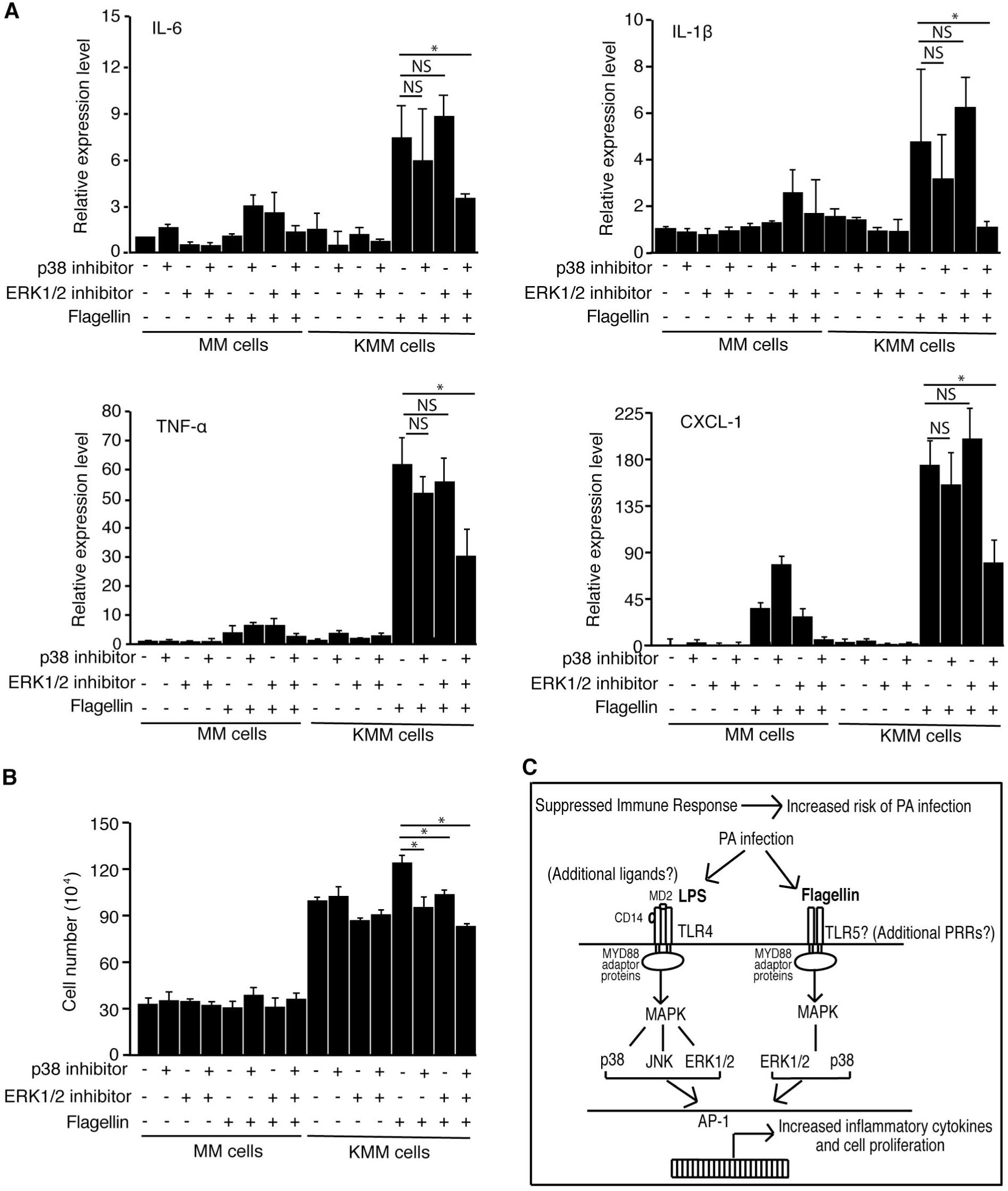
Simultaneous inhibition of p38 and ERK1/2 pathways decreases flagellin-induced inflammation and cell proliferation of KMM cells but has no obvious effect on MM cells. (A) MM and KMM cells were treated for 1 h with PBS, 10 μM p38 inhibitor (SB 203580), 10 μM ERK1/2 inhibitor (U0126), or a combination of 10 μM p38 inhibitor and 10 μM ERK1/2 inhibitor. Cells were then treated with PBS or 0.3 μg/mL flagellin for 1 h, and analyzed for cytokine by RT-qPCR. (B) MM and KMM cells were treated for 1 h with PBS, 10 μM p38 inhibitor (SB 203580), 10 μM ERK1/2 inhibitor (U0126), or a combination of 10 μM p38 inhibitor and 10 μM ERK1/2 inhibitor. Cells were then treated with 1 μg/mL flagellin and examined for cell number at day 3 post-treatment. (C) Schematic illustration summarizing PA ligands, and their receptors and activated downstream pathways, which might enhance inflammation and cell proliferation of KSHV-transformed cells during PA infection. Asterisk indicates P ≤ 0.05. NS denotes “not significant”.

We further assessed the roles of p38 and ERK1/2 MAPK pathways on flagellin-induced cell proliferation. While PA flagellin stimulated the proliferation of KMM cells by 20%, SB203580 or U0126 alone or in combination completely abolished this effect (Fig. 8B). SB203580 or U0126 alone or in combination had no significant effect on the proliferation of MM cells (Fig. 8B). These results indicated that both p38 and ERK1/2 MAPK pathways were essential for flagellin-stimulated proliferation of KMM cells.

## Discussion

It has been reported that the diversity of overall bacterial species is decreased while the number of pathogenic species is increased in the microbiomes of immunosuppressed patients compared to healthy individuals (39). The effects of pathogenic bacteria on cancer pathology have been studied extensively in numerous cancers (40). Pathogenic bacteria such as *Fusobacterium nucleatum, Helicobacter pylori*, and *Salmonella spp*. can exacerbate cancer pathogenesis by producing bacterial toxin and effector proteins that induce host cell damage and interfere with cell signaling pathways involved in cell proliferation (40). Additionally, bacterial metabolic products including short chain fatty acids can reactivate oncogenic viruses such as Epstein-Barr Virus and KSHV, resulting in dissemination of cancer cells (41). Our study specifically focuses on bacterial PAMPs, which activate TLRs, resulting in pro-survival pathway activation and increased cell proliferation (22).

In the current study, we investigated the role of PA in KSHV induction of chronic inflammation and KSHV-induced cell proliferation and cellular transformation. PA infection can occur in HIV-infected patients and can cause serious complications in immunocompetent hosts despite being a commensal bacterium (32). The results showed that PA stimulation enhanced cell proliferation, and the sizes and efficiency of colony formation in softagar of KSHV-transformed cells but had no effect on the uninfected primary cells. Mechanistically, PA stimulation increased the expression of inflammatory cytokines and activated multiple MAPK pathways. Importantly, besides LPS, we found that PA flagellin also contributed to the induction of inflammatory cytokines and cell proliferation of KSHV-transformed cells, and that LPS and flagellin differentially activated MAPK pathways but both induced similar profiles of inflammatory cytokines (Fig. 8C). Specifically, LPS activated p38, JNK, and ERK1/2 pathways while flagellin activated the p38 and ERK1/2 pathways despite both PAMPs inducing similar levels of inflammatory cytokines IL-6, IL-1β, TNF-α, and CXCL-1. Our results indicate that KSHV-transformed cells are more susceptible to PA-induced inflammation through PAMPs LPS and flagellin than the uninfected primary cells. It would be interesting to further confirm the roles of PA in KS development in AIDS-KS patients.

It has been shown that TLR activation leads to activation of downstream pro-survival pathways, resulting in increased cancer pathogenesis (42, 43). We previously showed that KSHV infection upregulated numerous TLRs including TLR4 and TLR5 (22). LPS and flagellin are highly immunogenic ligands for TLR4 and TLR5, respectively (33). Indeed, TLR4 was upregulated close to 50-fold in KSHV-transformed cells sensitizing the cells to TLR4 ligands (22). Activation of the TLR4 pathway by *E. coli* and LPS stimulated cell proliferation, cellular transformation, and tumorigenesis by increasing IL-6 expression to activate the STAT3 pathway (22). In this study, we demonstrate that both PA LPS and flagellin activates the MAPK pathways and that inhibition of MAPK pathways decreases expression of inflammatory cytokines, cell proliferation and cellular transformation (Fig. 8C).

We previously demonstrated that LPS stimulation increased cytokines IL-1β, IL-6, and IL-18 in KMM cells (22). In our current study, we showed that PA stimulation increased cytokines including IL-1β, IL-6, TNF-α, and CXCL-1 in KMM cells. Inflammatory cytokines such as TNF-α act synergistically with IL-6 to activate the STAT3 pathway (22, 44). Surprisingly, we did not see an increase in IL-18, despite previous studies reporting IL-18 upregulation correlating with PA infection (45, 46). It is possible that we had missed the peak time point for the induction of IL-18 as we focused on the effect of acute stimulation. As previously reported, we confirmed the induction of TNF-α and CXCL-1 by PA (35, 36, 47). Our results showed that both PA LPS and flagellin contributed to the induction of IL-1β, IL-6, TNF-α, and CXCL-1. It would be interesting to further assess the specific effects of LPS and flagellin in stimulating specific cytokines and their downstream inflammatory pathways in future studies.

The relationship between TLR activation and cancer cell proliferation has been extensively studied (42). Studies have demonstrated that LPS- and flagellin-induced inflammation increases cell proliferation in several cancer cell lines (13, 48, 49). Moreover, some studies suggest that LPS and flagellin blood levels may be correlated with a higher cancer risk (50, 51). However, contradictory studies show decreased cell proliferation in LPS- or flagellin-stimulated cells (52-54). In fact, bacteria and its PAMPS have been studied as antitumor agents (55, 56). It is worth mentioning that multiple factors might affect how cells respond to TLR activation. The differences in cell response to TLR activation may depend on the cellular location of the TLRs, as increased TLR expression in the cytoplasm can result in more chronic inflammation and cell proliferation. In contrast, TLR expression on the cell membrane may result in antitumor effects (57). The proportions of the types of TLRs expressed (TLR2 *vs*. TLR4, for example) may also affect in the cellular response (52, 58). There is no doubt that the tumor microenvironment, including the composition of immune cells, can substantially affect the response and proliferation of tumor cells (59). Activation of TLRs in immune cells can result in a Th1-response, causing apoptosis (59). Further elucidation of TLRs and their downstream pathways is essential for understanding the complex interactions between cell proliferation and inflammation.

Lastly, we focused on the effects of PA on MAPK activation, showing that PA stimulation of KMM cells resulted in the activation of p38, ERK1/2, and JNK pathways. PA flagellin contributed to cell proliferation by activating the p38 and ERK1/2 pathways and inhibition of the p38 and ERK1/2 pathway abrogated the enhanced cell proliferation in flagellin-stimulated KMM cells (Fig. 8C). The MAPK pathways play an important role in KSHV biology and likely in KS pathology. Primary infection of KSHV results in activation of p38, ERK1/2, and JNK activation and inhibition of the MAPK pathways can reduce KSHV infectivity and induction of IL-6 (24, 60). The MAPK pathways also mediate KSHV reactivation (61). Our results identify the important role of PA-induced inflammation in cell proliferation and cellular transformation while providing further evidence in the therapeutic value of inhibiting the MAPK pathways in KS patients.

Overall, our study utilizes a clinically relevant, opportunistic bacterium to study the effect of inflammation on KSHV-transformed cells. We report that PA stimulation increases cell proliferation and cellular transformation in KSHV-transformed cells, while having no significant effect on MM cells. These results emphasize that KSHV cellular transformation results in enhanced sensitivity to external stimuli, which may further increase cell proliferation and cellular transformation. As KS and HIV/AIDS patients are at an increased risk for opportunistic infection, it is critical to understand the effects of bacteria on KS pathogenesis. Our results indicate that elimination of certain bacterial infections identified to promote inflammation may have preventive value for KSHV-infected AIDS patients who are at a high risk for developing KS as well as therapeutic value for AIDS-KS patients. Moreover, our study further dissects the specific PA PAMPs that contribute to cell proliferation, demonstrating that both LPS and flagellin can induce inflammation in KSHV-transformed cells. Besides LPS and flagellin, PA consists of many additional PAMPs that can induce inflammation, such as peptidoglycans and lipoproteins (62, 63). Analyzing the effects of these additional PAMPs on cell proliferation, by exploring the specific pathways and mechanisms of inflammatory induction, may clarify the correlative, additive or synergistic effects as well as independent effects of these other PAMPS in inflammation process in general as well as for the inflammation and KS pathogenesis specifically.

## Materials and Methods

### Cell Culture

Early passages (<20) of MM, KMM, BJAB, and KSHV-BJAB cells were grown as previously described (34, 64). All cell lines were routinely tested for mycoplasma contamination using LookOut Mycoplasma qPCR Detection Kit (Sigma, MP0035-1KT).

### Reagents

Purified flagellin from PA (tlrl-pafla, InvivoGen) and Ultrapure LPS from *E. coli* K12 (tlrl-peklps, InvivoGen) were resuspended in water. CLI095 (tlrl-cli095, Thermo Fisher Scientific), SB203850 (NC9041893, Fisher), and U0126 (19-147, Sigma) were resuspended in DMSO.

### Bacterial preparation

Three PA strains were used for this study: PA (ATCC^®^ 15442™), PAO1Δ*fliC* (JJH325), and PAO1. PAO1Δ*fliC* and PAO1 were kindly provided by Dr. Jennifer Bomberger (University of Pittsburgh), Dr. Joe Harrison (University of Calgary), and Dr. Matthew Parsek (University of Washington). PA strains were grown in LB broth (Sigma Aldrich) overnight until an OD value of 1.1. The culture was then washed three times in PBS by centrifuging at 3,200 g for 10 min and then diluted with PBS to the specified concentrations (CFUs/mL) for experiments. To confirm accuracy of the bacterial concentration, the culture was serially diluted, grown on agar plates overnight, and analyzed for number of colonies.

### Cell proliferation assay

MM/KMM cells and BJAB/KSHV-BJAB cells were plated at a density of 20,000 and 10,000 cells/well (respectively) in 24-well plates for 16 h, treated with the indicated reagents, and counted using a Malassez chamber.

### Softagar assay

Colony formation in softagar was carried out as previously described (34).

### RNA extraction and qRT-PCR

Total RNAs were extracted with the TRI Reagent (Sigma, T9424). Reverse transcription (RT) was performed with 1 μg of total RNA using SsoAdvanced™ Universal SYBR® Green Supermix (Bio-Rad, 172-5272). cDNAs diluted 2 times were examined by qPCR with the KAPA SYBR Fast qPCR Kit (Kapa Biosystems, K4602) using specific primers for β-actin, IL-6, IL-1β, IL-18, TNF-α and CXCL-1. β-actin gene was used for loading calibration. All the sequences of primers used for qRT-PCR are listed in Table 1.

**Table 1.**
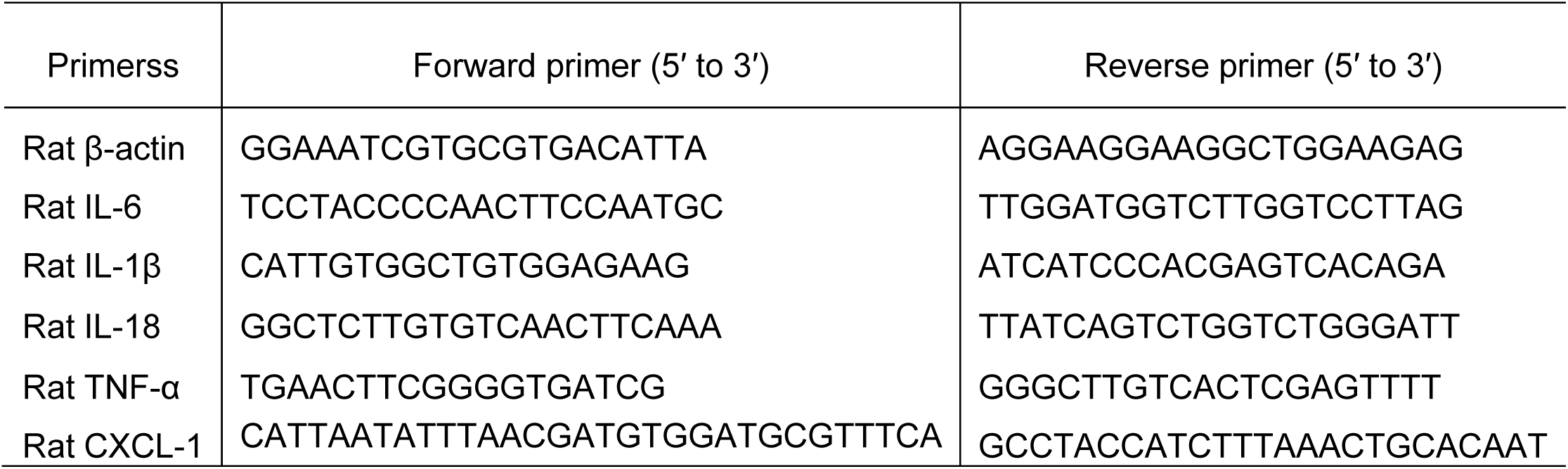
Sequences of primers used for RT-qPCR.

### Western-blotting analysis

Western blot analysis was performed as previously described (65). Primary antibodies included mouse mAbs to β-tubulin, and rabbit polyclonal antibodies to GAPDH (5174S, Cell Signaling Technology), JNK (9252S, Cell Signaling Technology), phospho-JNK (4668S, Cell Signaling Techology), p38 (8690S, Cell Signaling Technology), phospho-p38 (4511S, Cell Signaling Technology), ERK1/2 (4695S, Cell Signaling Technology), phospho-ERK1/2 (4370S, Cell Signaling Technology).

### Statistical analysis

Results were expressed as mean ± SE from at least three independent experiments. Statistical analysis was performed using two-tailed t-test and P ≤ 0.05 was considered significant.

## Acknowledgments

We thank Dr. Jennifer Bomberger of the University of Pittsburgh, Dr. Joe Harrison of the University of Calgary, and Dr. Matthew Parsek of the University of Washington for providing PA strains. This work was supported by grants from the National Institute of Health (CA096512, CA124332, CA132637, CA213275, DE025465 and CA197153 to S-J Gao). This work was supported in part by award P30CA047904. We thank members of Dr. Gao’s laboratory for technical assistances and helpful discussions.

